# Preferential synthesis of very long chain polyunsaturated fatty acids in *eutreptiella* sp. (eugelnozoa) revealed by chromatographic and transcriptomic analyses

**DOI:** 10.1101/2020.05.14.097162

**Authors:** Rita C. Kuo, Huan Zhang, James D. Stuart, Anthony A. Provatas, Linda Hannick, Senjie Lin

**Affiliations:** Department of Marine Sciences, University of Connecticut, Groton, CT 06340, USA; Department of Chemistry, University of Connecticut, Storrs, CT 06269, USA; Center of Environmental Sciences and Engineering, University of Connecticut, Storrs, CT 06269, USA; SAIC-Frederick, Inc., Frederick National Laboratory for Cancer Research, Rockville, MD 20852, USA

**Keywords:** *Eutreptiella*, lipid biosynthesis, nutrient limitation, transcriptomic, unsaturated fatty acids

## Abstract

Algal lipids are important fuel storage molecules in algae and a currency for energy transfer in the marine food chain as well as materials for biofuel production, but their production and regulation are not well understood in many species including the common coastal phytoplankton *Eutreptiella* spp. Here, using gas chromatography-tandem mass spectrometry (GC/MS/MS), we discovered 24 types of fatty acids (FAs) in *Eutreptiella* sp. with a relatively high proportion of long chain unsaturated FAs. The abundances of C16, C18 and saturated FAs decreased when phosphate in the culture medium was depleted. Among the 24 FAs, docosahexaenoic acid (22:6) and eicosapentaenoic acid (20:5) were the most abundant, suggesting that *Eutreptiella* sp. preferentially invests in the synthesis of very long chain polyunsaturated fatty acids (VLCPFA). Further transcriptomic analysis revealed that *Eutreptiella* sp. likely synthesizes VLCPFA via Δ8 pathway and uses type I and II fatty acid synthases. Using RT-qPCR, we found that some of the lipid production genes, such as β-ketoacyl-ACP reductase, fatty acid desaturase, acetyl-CoA carboxylase, acyl carrier protein, Δ8 desaturase, and Acyl-ACP thioesterase, were more actively expressed during light period. Besides, two carbon-fixation genes were more highly expressed in the high lipid illuminated cultures, suggesting a linkage between photosynthesis and lipid production.

## Introduction

Lipids are important fuel storage molecules in algae as well as other organisms, and as algae are the base of the aquatic ecosystem, lipids are an important currency of energy transfer in the aquatic food chain (Jonasdottir 2019). Lipid content also represents the quality of an alga as materials for biofuel production (Sajjadi et al. 2018). Within algae, lipids are essential for structural constituents of cellular membranes in all organisms and protection of delicate internal organs and hormones in animals (Singh 2002). In the marine ecosystem, fatty acids are a fundamental energy source for growth and cell membrane fluidity (especially in cold water) in animals such as zooplankton and fish (*e.g.* Arzel et al. 1994, Cossins et al. 1977, Klein Breteler 2004, Rainuzzo et al. 1997, Tang and Taal 2005), which consume phytoplankton as food. To gain better understanding on the energy flow in marine food chains, it is important to study fatty acid and lipid biosynthesis pathways in dominant phytoplankton (prey) species. As the type and abundance of lipids in the cells also determine the potential of algae as source of biofuel, understanding the regulatory pathways of lipid production in algae is also important for evaluating or genetically enhancing algal species for biofuel production.

In land plants, photosynthesis produces small precursors for lipid production in chloroplasts. The small precursors are later converted into long chain fatty acids via two enzyme systems, acetyl-CoA carboxylase (ACCase) and fatty acid synthase (FAS). ACCase is essential for lipid metabolism as it catalyzes the first step of fatty acid synthesis and is found in both bacteria and eukaryotes (Cronan and Waldrop 2002, Harwood 1988). FAS consists of 6 enzyme activities and an acyl-carrier protein (CA) to catalyze a 2 carbon elongation process (Bloch and Vance 1977, Nelson and Cox 2008). It produces palmitate and stearate and these fatty acids are subject to elongation, desaturation or further modification by elongase or desaturase in chloroplasts (Gunstone et al. 2007). There are two major types of fatty acid synthases. Fatty acid synthase I (FASI), found in vertebrates and fungi, consists of a multi-enzyme complex contained in a single polypeptide chain (Nelson and Cox 2008, Schweizer and Hofmann 2004). In contrast, fatty acid synthase II (FASII), found in plants and bacteria, contains discrete enzymes (Nelson and Cox 2008, White et al. 2005). Among euglenoid algae, *Euglena gracilis* composes both FASI and plastid FASII for *de novo* fatty acid synthesis (Delo et al. 1971, Worsham et al. 1993). In addition to FASII, Goldberg and Bloch (1972) reported another ACP-dependent fatty acid synthase (FASIII) that elongates acyl-CoA derivatives from C10 to C18 to longer chain ACP thioesters in plastids.

Similar to land plants, microalgae initiate fatty acid synthesis in chloroplasts (Guschina and Hardwood, 2006) and lipid biosynthesis is probably linked directly to photosynthesis. Studies have shown that the activities of ACCase are related to the light reactions of photosynthesis, as NADPH and ATP produced in the photochemical reaction are required for the synthesis of palmitic acid (Sasaki et al. 1997). Furthermore, fatty acid synthesis in photosynthetic organisms obviously relies on carbon fixation for carbon precursors for fatty acid synthesis (Bao et al. 2000). The synthesized fatty acids are in turn used for triacylglycerol (TAG) synthesis (Thelen and Ohlrogge 2002). In addition, dihydroxyacetone phosphate, the precursor for glycerol 3-phosphate needed in the synthesis of lipid TAG, can be produced from Calvin-Benson Cycle (Nelson and Cox 2008).

It is well understood that lipid production in microalgae is influenced by nutrient and other environmental conditions (e.g. Gouveia and Oliveira 2009, Griffiths and Harrison 2009, Hu et al. 2008, Illman et al. 2000, Liu et al. 2008). Nutrient limitation can enhance the production of lipid in algal cells. For example, the lipid content of some *Chlorella* species (*e.g. C. emersonil and C. pyrenoldosa*) can increase to more than 60% of the dry weight under nitrogen-deprived conditions (Griffiths and Harrison 2009, Illman et al. 2000). However, our knowledge of the genetic regulatory mechanisms of lipid biosynthesis in marine microalgae is limited and fragmentary (Guchina and Harwood 2006, Hu et al., 2008), particularly for euglenids such as *Eutreptiella* spp.

*Eutreptiella* is a genus of photosynthetic euglenoid, which have excellent nutritive value (*i.e.* high vitamin and lipid content, Takeyama et al., 1996; Yamane et al., 2001) and are rather abundant in some marine ecosystems (Henriksen et al. 2002). Some *Eutreptiella* spp. can seasonally be a dominant group of phytoplankton (Álvarez-Góngora and Herrera-Silveira 2006, Bates and Strain 2006, Olli et al. 1996, Rodríguez-Graña et al. 2008, Seong et al. 2006), forming blooms in nutrient-rich coastal or brackish waters (Anderson et al. 2000, Lindholm 1993, Olli et al. 1996, Stonik and Selina 200, Stonik 2007). In some areas, *Eutreptiella braarudii* alone can make up to 46% of the phytoplankton population (Stonik, 2007). As primary producers, these algae are important in energy flow and nutrient cycling in the coastal marine ecosystem. Meanwhile, as solar energy converters, these algae are potentially candidates of biofuel species.

In this study, we used 454 high-throughput sequencing to study transcriptomic profiles and gas chromatography-tandem mass spectrometry (GC/MS/MS) to identify fatty acids produced under phosphate-depleted and phosphate-replete conditions, and conducted reverse-transcription qPCR (RT-PCR) to quantify expression of several key enzyme coding genes to investigate molecular mechanisms that regulate lipid biosynthesis in *Eutreptiella* sp.

## Materials and Methods

### Culture preparation and RNA isolation

Two sets of cultures were maintained in phosphate-depleted medium (to produce high-lipid culture) and phosphate-replete f/2 medium (to produce low-lipid culture); cells were harvested, total RNA and mRNA were isolated as reported previously (Kuo et al. 2013)

### Lipid measurement

Nile Red (9-diethylamino-5H-benzo-*α*-phenoxazine-5-one, a lipid-soluble fluorescent probe) staining was used to measure relative abundance of neutral lipids as reported (Lee et al., 1998, Hu *et al*., 2008). Forty μl of Nile Red solution in acetone (250 mg/l) were added to 10 ml of algal suspension at room temperature for 10 min. A spectrophotometer (HITACHI, Tokyo, Japan) was then used to measure the fluorescence with excitation at the wavelength of 490 nm and emission in the wavelength band of 580 to 590 nm, as reported in Kuo and Lin (2012).

### Lipid extraction and fatty acid identification by gas chromatography-tandem mass spectrometry (GC/MS/MS)

A gravimetric method was applied to determine actual lipid contents by using chloroform-methanol method according to Bligh and Dyer (1959). In short, lipids were extracted from freeze-dried cells with chloroform/methanol (1:2, v/v) and the residue was extracted one more time with 1:1 chloroform/methanol. The chloroform layer was collected and evaporated under a gentle flow of nitrogen gas. Derivatization for GC/MS/MS analysis followed the method summarized by (Carvalho and Malcata 2005). Briefly, samples were dissolved in 1 ml of a freshly prepared mixture of dry acetyl chloride and methanol, at a ratio of 5:100 (v/v), and kept at 100 °C under pure nitrogen for 1 h. After cooling, 1 ml of hexane wad added and mixed by vortexing. Purification of the solution was achieved by adding 1 ml of saturated sodium chloride solution. The prepared solution of fatty acid methyl esters was filtered using 0.45 μm Millipore filter. Analysis of the solution was performed on a Waters Quattro Micro GC/MS/MS system equipped with a Rxi-5Sil MS column (30 m x 0.25 mm, 0.25 μm film thickness, Restek Co., PA). Helium was used as the carrier gas with a flow rate of 1 ml/min. Initial oven temperature was 70 °C for 1 min and the temperature gradient was 5 °C/min from 70 °C to 270 °C with no holding time. Sample was introduced in splitless injection mode. Injector temperature was 270 °C and purged for 1 min with a purge flow of 25 ml/min. Total run time was 41 min. The MS source temperature was set at 250 °C and the GC interface was 275 °C. A fatty acids methyl ester mixture (FAMQ-005 FAME Reference Standard, AccuStandard Inc., CT, USA) was used as external standards. The mass spectrum and the corresponding retention time of each component were exported to NIST MS Search 2.0 software to identify compounds by means of library and standard comparisons.

### 454 sequencing of the SL-based transcriptomes

mRNA samples isolated as described above from the four culture conditions were used to synthesize cDNA, which was subsequently used for 454 sequencing as reported previously (Kuo et al. 2013). Briefly, a modified random oligo named 454AT_7_N_9_ (5’-CGTATCGCCTCCCTCGCGCCATCAGTAATACGACTCACTATAGGGAGNNNNNNNNN-3’, where N is any of the 4 nucleotides) was used to synthesize the 1st strand cDNAs. The cDNAs were then used as templates for PCR amplification of the 5’-end of the cDNAs of the *trans-*spliced transcripts using ExTaq with a *Eutreptiella* spliced leader-based primer 454BEutSL, 5’-GAGACTATGCGCCTTGCCAGCCCGCTCAGACACTTTCTGAGTGTCTATTTCTTTTCG-3’), paired with 454AT_7_ (5’-CGTATCGCCTCCCTCGCGCCATCAGTAATACGACTCACTATAGGGAG-3’). After PCR we selected the amplicons from agarose gel in the size range of 300-700 bp as the template for emulsion PCR (emPCR) using GS Titanium SV emPCR Kit. Sequencing was carried out using GS Titanium Sequencing Kit on the GS FLX System at the Center for Applied Genetics and Technology, University of Connecticut.

### Sequence processing and annotation

Sequence reads were processed and analyzed as recently reported (Kuo et al. 2013). In short, raw sequencing reads from all of the 4 samples were pooled together and trimmed using CLC Genomics Workbench (CLC Bio, Aarhus, Demark). After quality and primer trimming, sequences shorter than 150 nt were discarded. In order to filter off 454 sequencing errors and create reduced-redundancy sequence dataset, USEARCH (Edgar 2010) was used for sequence clustering. The resulting unique transcripts were then annotated using Blast2Go V.2.5.0 (Götz et al. 2008) against NCBI’s non-redundant (nr) database using BLASTx algorithm (Altschul et al. 1990), with a cut-off-E-value ≤ 10^−3^.

### Reverse-transcription quantitative PCR (RT-qPCR)

In order to further investigate the expression patterns of genes potentially regulating lipid production in *Eutreptiella*, some of the genes identified from the transcriptomic data as related to carbon fixation and lipid synthesis were selected and further analyzed by RT-qPCR. First strand cDNA libraries were prepared by the methods mentioned earlier. It is suggested that at least two reference genes are needed for proper normalization of gene expression levels (Guo and Ki 2012, Vandesompele et al. 2002). Tubulin, actin, and elongation factor 1α were selected as candidates, as they had been shown to be good reference genes in some microalgae and plants (Guo and Ki 2012, Le Bail et al. 2008). We used the geometric average of the expression levels of these reference genes to normalize the measured expression levels of target genes as reported (Vandesompele et al. 2002). For each gene investigated, two sets of primers were designed (Table 3). To prepare standards, the primer combination that specifies longer gene fragment was used to PCR amplify each target gene from *Eutreptiella* cDNA. The amplicons of the target genes were checked with gel electrophoresis to assure the absence of primer dimmers and then purified using DNA Clean & Concentrate™ column (Zymo Research, Orange, CA). The purified DNA was serially diluted to 10^2^, 10^3^, 10^4^, 10^5^, and 10^6^ gene copies to generate standard curves to analyze the amplification efficiency and primer specificity for every primer pair. qPCR was performed using the iCycler iQ™ real-time PCR Detection System (Bio-Rad Laboratories, Inc., Hercules, CA) with SYBR Green supermix. The qPCR reactions included a single denaturation cycle of 95 °C for 3 min, 40 cycles of 95 °C for 20 sec, 58 °C for 30 sec, and 72 °C for 15 sec, followed by a melt curve analysis from 55 to 100 °C.

**Table 1.**
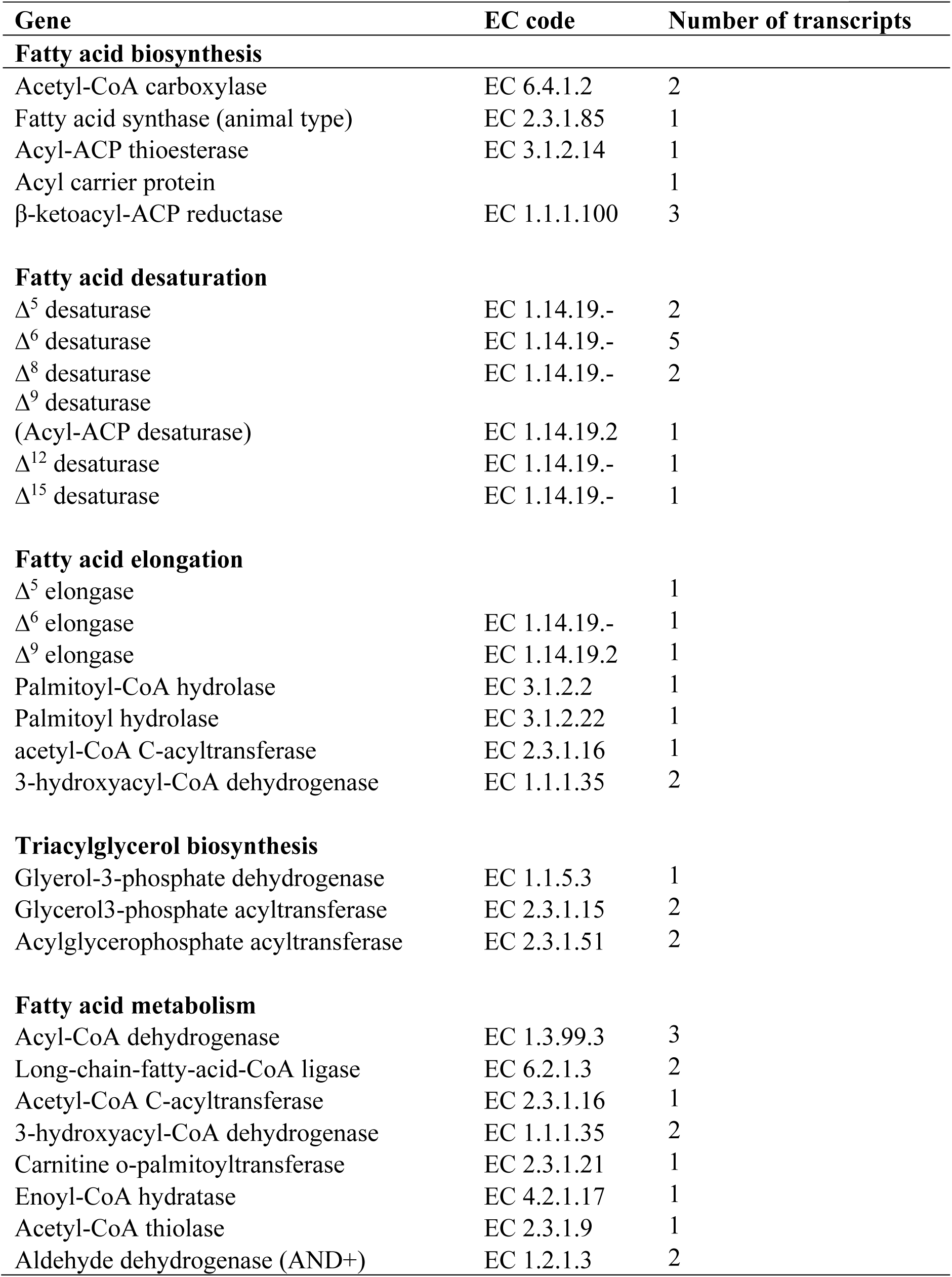
Genes potentially involved in fatty lipid biosynthesis in *Eutreptiella* sp. Sequences and BLAST results were listed in the supporting file “lipid_biosynthesis_genes”.

**Table 2.** Types and relative abundance of fatty acids detected in *Eutreptiella* sp.

| Peak ID | Retention time | Systematic name | Abbreviation | Relative abundance (%)F/2* | Relative abundance (%)F/2-P* |
| --- | --- | --- | --- | --- | --- |
| 1 | 26.19 | 4,7,10,13-Hexadecatetraenoic acid | 16:4(n-3) | 98.04 | 11.44 |
| 2 | 26.37 | 7,10-Hexadecadienoic acid | 16:2(n-6) | 2.59 | 0.08 |
| 3 | 26.47 | 7,10,13-Hexadecatrienoic acid | 16:3(n-3) | 12.63 | 0.54 |
| 4 | 26.62 | 7-Hexadecenoid acid | 16:1(n-9) | 11.89 | 0.27 |
| 5 | 26.97 | 9-Hexadecenoid acid | 16:1(n-7) | 13.54 | 0.79 |
| 6 | 27.1 | Hexadecanoic acid | 16:0 | 90.45 | 11.28 |
| 7 | 29.11 | 6,9,12-Octadecatrienoic acid | 18:3(n-6) | 3.90 | 0.13 |
| 8 | 30.04 | 6,9,12,15-Octadecatetraenoic acid | 18:4(n-3) | 41.00 | 12.40 |
| 9 | 30.23 | 9,12-Octadecadienoic acid | 18:2(n-6) | 3.20 | 0.53 |
| 10 | 30.36 | 9,12,15-Octadecatrienoic acid | 18:3(n-3) | 72.63 | 23.58 |
| 11 | 30.47 | 9-Octadecenoic acid | 18:1(n-9) | 7.97 | 0.58 |
| 12 | 30.85 | Octadecanoic acid | 18:0 | 8.43 | 2.12 |
| 13 | 33.1 | 5,8,11,14-Eicosatetraenoic acid | 20:4(n-6) | 6.44 | 1.88 |
| 14 | 33.22 | 5,8,11,14,17-Eicosapentaenoic acid | 20:5(n-3) | 78.69 | 40.02 |
| 15 | 33.53 | 8,11,14,17-Eicosatetraenoic acid | 20:4(n-3) | 8.39 | 4.02 |
| 16 | 33.77 | 8,11-Eicosadienoic acid | 20:2(n-9) | 1.13 | 0.71 |
| 17 | 33.87 | 11,14,17-Eicosatrienoic acid | 20:3(n-3) | 3.16 | 1.02 |
| 18 | 36.17 | 4,7,10,13,16-Docosapentaenoic acid | 22:5(n-6) | 68.78 | 54.93 |
| 19 | 36.3 | 4,7,10,13,16,19-Docosahexaenoic acid | 22:6(n-3) | 100.00 | 100.00 |
| 20 | 36.4 | 7,10,13,16-Docosatetraenoic acid | 22:4(n-6) | 43.11 | 47.18 |
| 21 | 36.49 | 7,10,13,16,19-Docosapentaenoic acid | 22:5(n-3) | 3.56 | 6.03 |
| 22 | 37.54 | Docosanoic acid | 22:0 | 5.94 | 1.01 |
| 23 | 38.79 | 6,9,12,15,18,21-Tetracosahexaenoic acid | 24:6 (n-3) | 5.32 | 0.85 |
| 24 | 40.14 | 15-Tetracosenoic acid | 24:1(n-9) | 0.46 | - |

\*The numbers of relative abundance were obtained by dividing the peak area of the fatty acid to the peak area of DHA; F/2 and F/2-P depict phosphate-replete and phosphate-depleted F/2 medium, respectively.

**Table 3.**
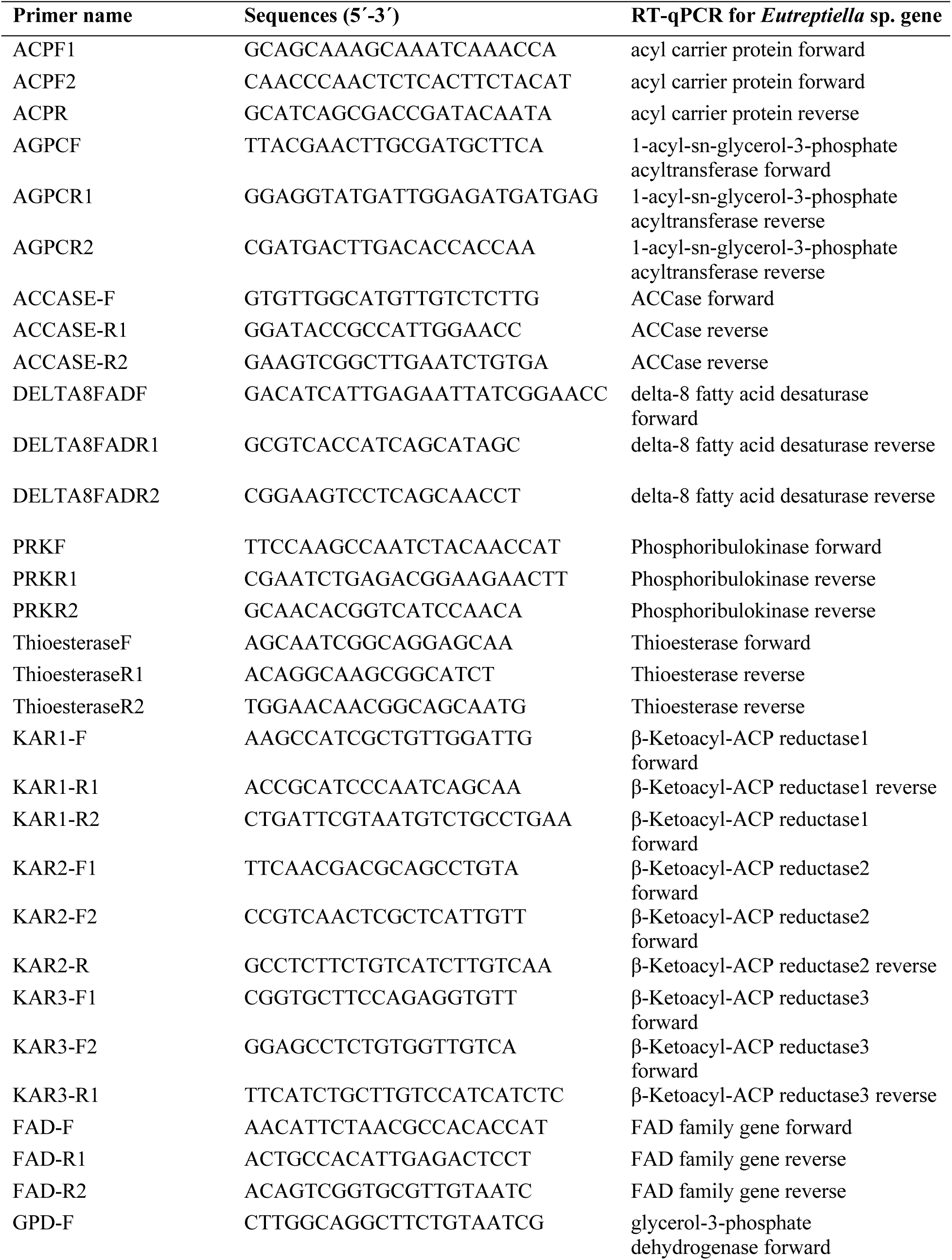
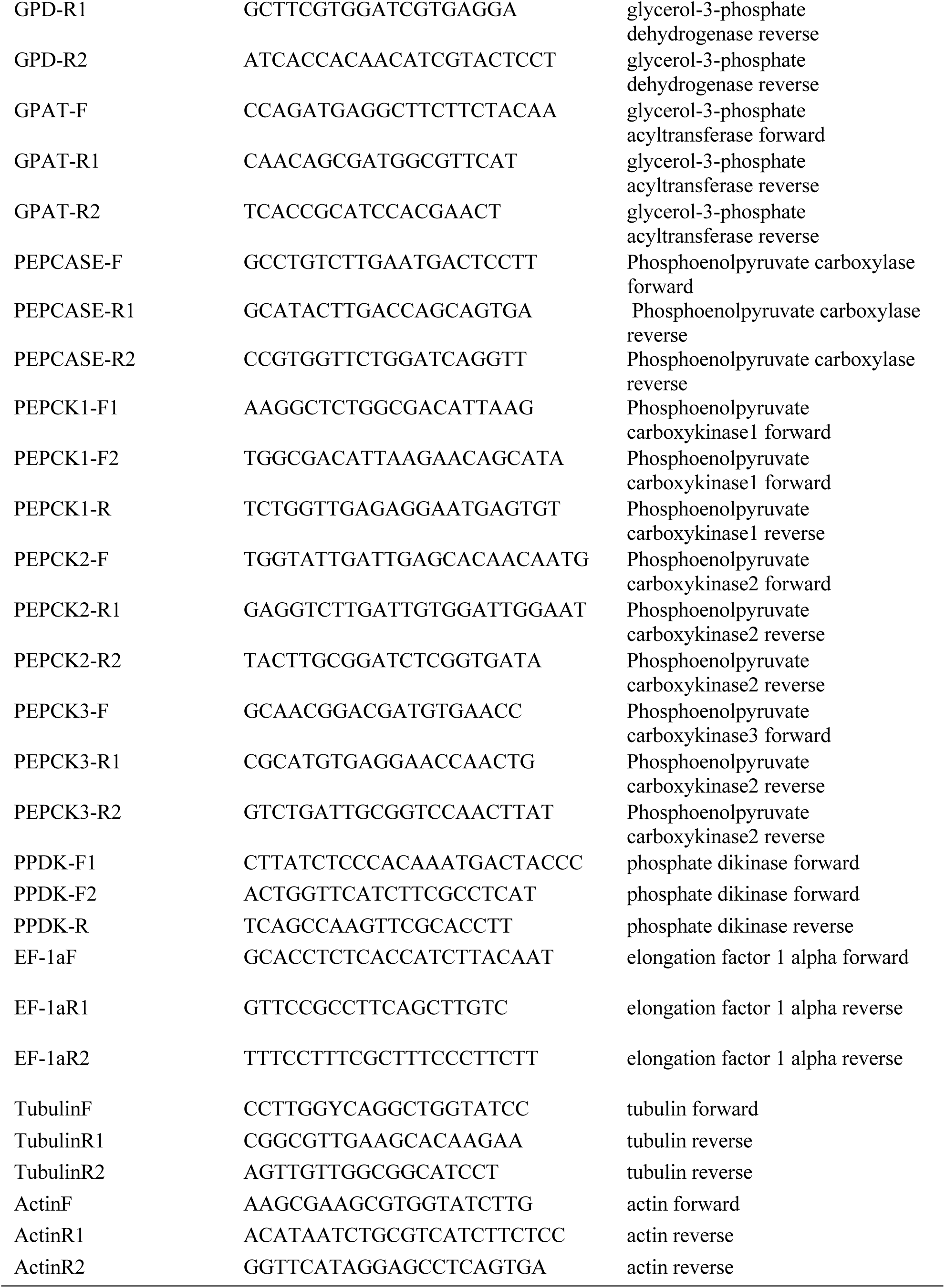
Primers used in reverse-transcription quantitative PCR (RT-qPCR) analysis in this study.

## Results

### Physiological conditions of the cultures

Two sets of cultures were grown under phosphate-depleted (to produce high-lipid culture) and phosphate-replete (to produce low-lipid culture) conditions. On the 11th day, the phosphate-replete cultures were in exponential growth phase whereas phosphate-depleted cultures already reached stationary phase. When normalized to per cell basis, the lipid contents of the cultures grown in phosphate-depleted medium were about 2-fold higher than that of the cultures grown in phosphate-replete medium (Figure 1). This observed difference in lipid content prompted cell harvesting from both cultures in the light and dark periods, which yielded four samples referred to as high-lipid-light, high-lipid-dark, low-lipid-light, and low-lipid-dark cultures.

**Figure 1.**
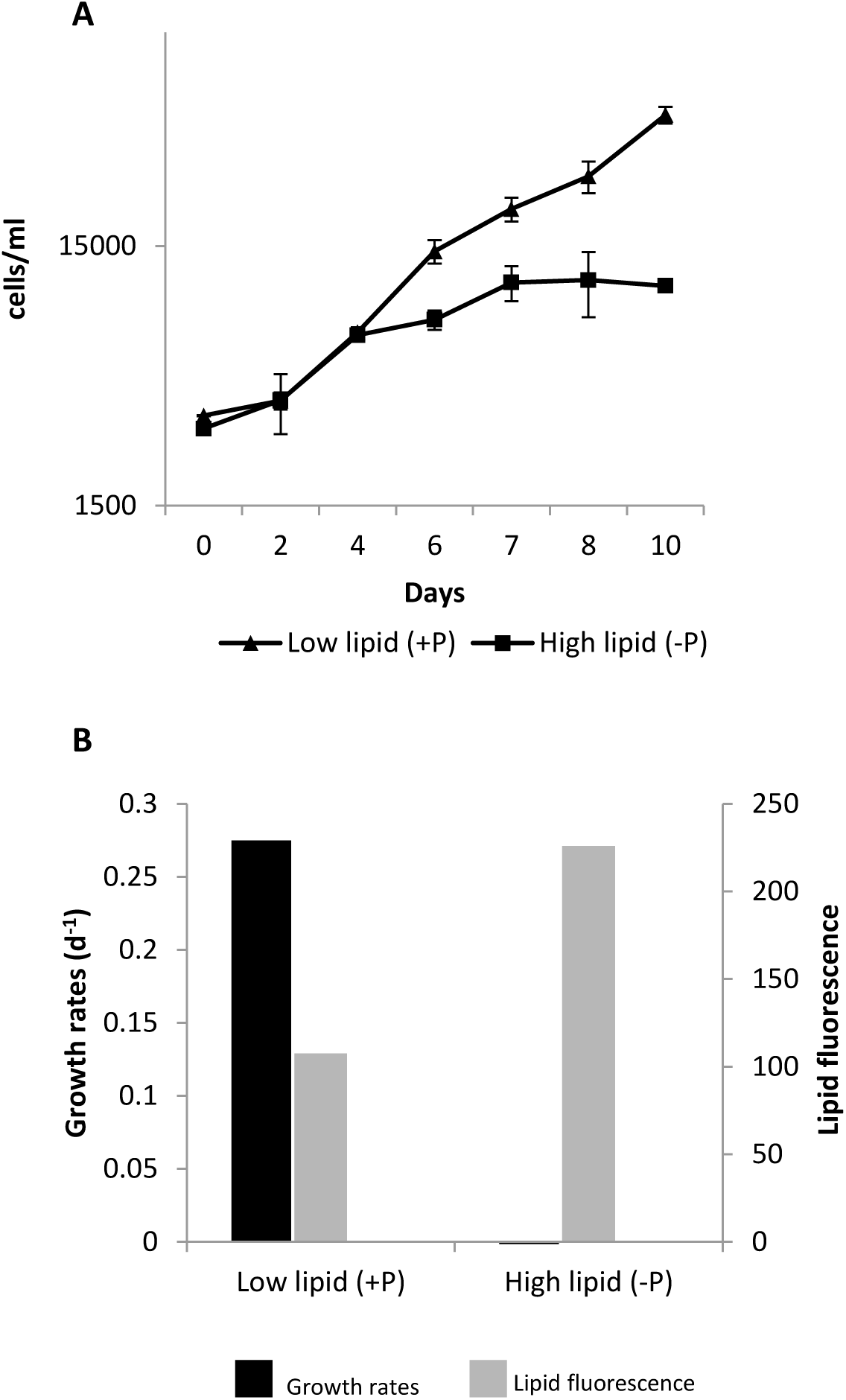
Growth and lipid production of *Eutreptiella* cultures under phosphate-replete and phosphate-depleted conditions. (A) Growth curves of *Eutreptiella* cultures. (B) Growth rates and lipid fluorescence levels at the time when cultures were harvested.

### Comparison of gene diversity among samples

The four samples were subjected to RNA extraction, cDNA library construction, and 454 sequencing (Kuo et al. 2013). The Venn diagram in Figure 2 summarizes the numbers of unique genes, total genes and shared genes among the 4 cDNA libraries. The analysis revealed that each library contained different numbers of total genes as well as different unique genes that were not shared by any other libraries. Between the two light cultures, the high-lipid cultures had more unique genes and total genes. Between the cultures in the same lipid content categories, cultures harvested under the light condition displayed more unique genes (i.e. light-specific) and total genes. Overall, high-lipid-light cultures expressed the highest number of unique and total genes among the 4 samples, suggesting that a more complex system was involved in lipid production and light reaction in *Eutreptiella* sp.

**Figure 2.**
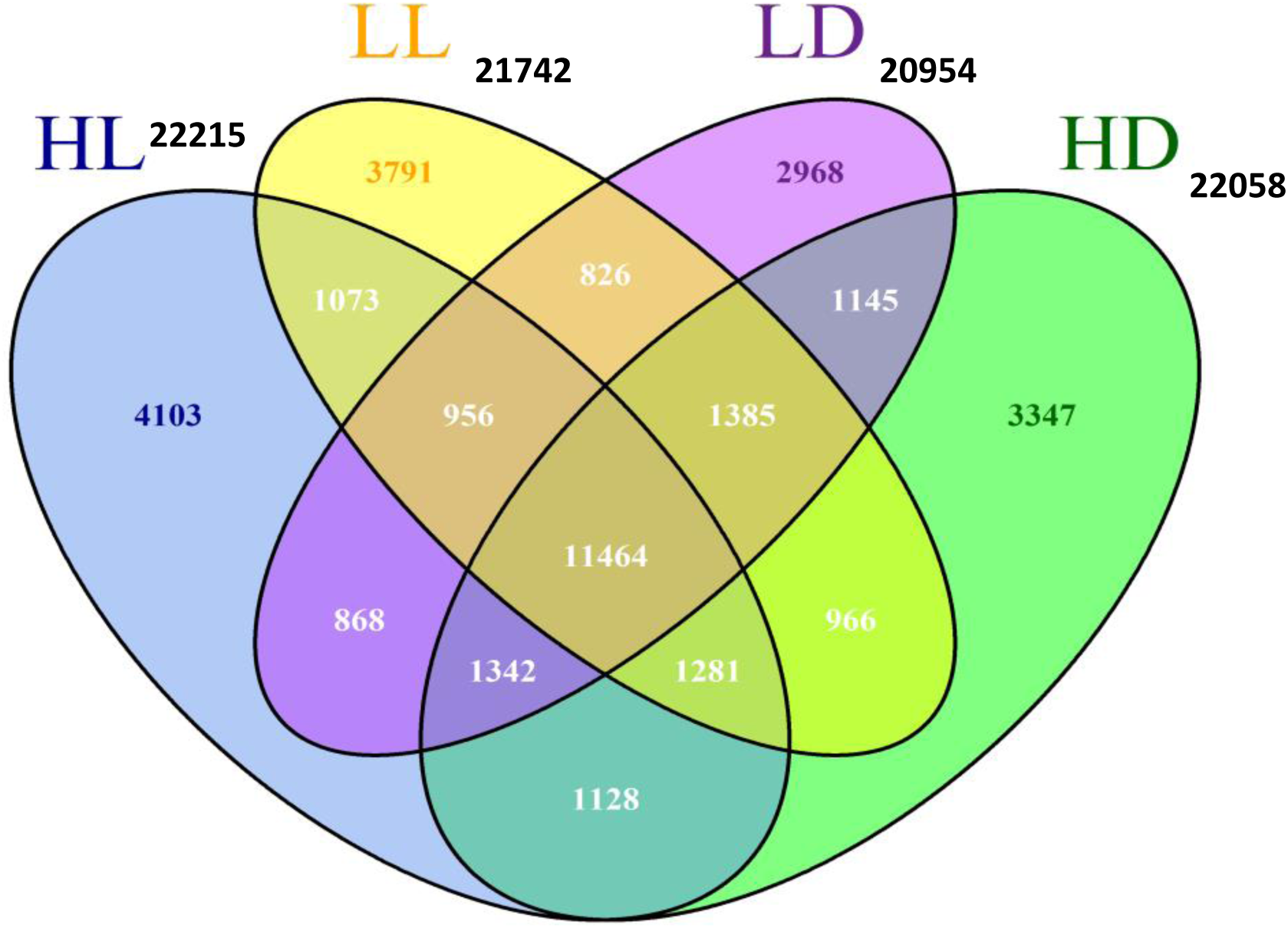
Common and unique expressed genes between different culture conditions. HL: high-lipid-light cultures. HD: high-lipid-dark cultures. LL: low-lipid-light cultures. LD: low-lipid-dark cultures. Numbers outside the diagram represent total number of genes found in each library.

### Fatty acid synthesis

Based on the functional annotation, we have identified genes encoding key enzymes involved in the lipid biosynthesis and catabolism (Table 1). Fatty acid biosynthesis from acetyl-CoA involved in two enzymes: ACCase and FAS. Eukaryotic type ACCase (72% identical to ACCase of *Thalassiosira pseudonana*; GenBank accession number: XP_002296083) and type I FAS (67% identical to FASI of *Caenorhabditis elegans*; NP_492417) were found in our dataset. Figure 3 illustrates a potential lipid biosynthesis pathway of *Eutreptiella* sp. ACCase catalyzes carboxylation of acetyl-CoA to produce malonyl-CoA, which is catalyzed to malonyl-ACP by malonyl-CoA transacylase. Next, long-chain saturated fatty acids are synthesized by type I fatty acid synthase, which have 7 active sites within a single large polypeptide complex (Nelson and Cox, 2008). The active enzyme domains include β-ketoacyl-ACP synthase (KAS), malonyl/acetyl-CoA-ACP transferase (MAT), β –hydroxyacyl-ACP dehydratase (HAD), enoyl-ACP reductase (EAR), β –ketoacyl-ACP reductase (KAR), and Acyl-ACP thioesterase (TE). Synthesized palmitate (C16:0) and stearate (C18:0) via the activity of FASI are further subjected to elongation, desaturation, or other modifications (see next section for more details). Besides FASI, individual genes encoding β –ketoacyl-ACP reductase and Acyl-ACP thioesterase related to type II fatty acid synthase (FASII) were also identified (Table 1). Malonyl-CoA transacylase, which functions in the fatty acid synthesis pathway in other plants, was missing in our dataset.

**Figure 3.**
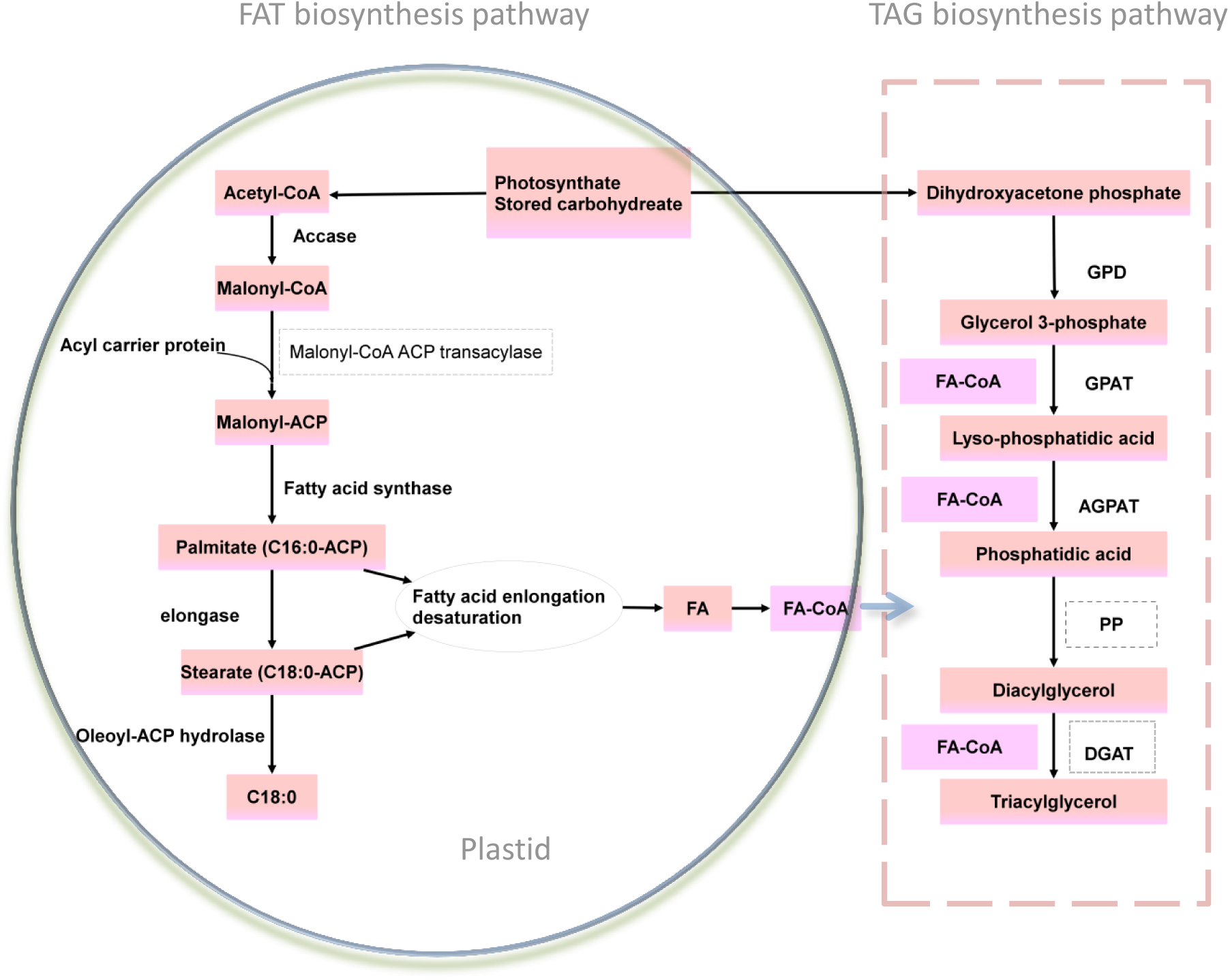
Putative lipid biosynthesis pathway in *Eutreptiella* sp. inferred from our transcriptomic data. GPD: glycerol 3-phosphate dehydrogenase (EC: 1.1.1.53); GPAT: glycerol-3-phosphate acyltransferase (EC: 2.3.1.15); AGPAT: acylglycerophosphate acyltransferase (EC: 2.3.1.51); PP: phosphatidate phosphatase (EC: 3.1.3.4); DGAT: diacylglycerol acyltransferase (EC: 2.3.1.20); FA: fatty acids; FA-CoA: fatty acyl-CoAs. FA: fatty acids. Dashed boxes indicate enzymes that were not found in our dataset. The process of FA elongation and desaturation is shown in Figure 5.

### Fatty acid identification and biosynthesis pathways

The stepwise desaturation and elongation of C18 acid lead to the extension of fatty acids and synthesis of unsaturated fatty acids (Nelson and Cox, 2008). The major fatty acids found in *Eutreptiella* sp. were the very long chain polyunsaturated fatty acids, docosahexaenoic acid (DHA, 22:6n-3). Various desaturases and elongases were found in our study (Table 1), including Δ5 desaturase, Δ6 desaturase, Δ8 desaturase, Δ9 desaturase, Δ12 desaturase and Δ15 desaturase. The elongases were: Δ5 elongase, Δ6 elongase, and Δ9 elongase.

In order to link the identified genes to their expected roles in *Eutreptiella* sp., the fatty acid (FA) profiles of *Eutreptiella* sp. were analyzed by screening C16-C24 fatty acids via GC/MS/MS. We identified 24 fatty acids (including C16, C20, C22, and C24 fatty acids) from the cultures of *Eutreptiella* sp. (Figure 4 and Table 2). In both of the phosphate-depleted and phosphate-replete cultures, *Eutreptiella* sp. was relatively rich in polyunsaturated fatty acids (PUFAs), especially very long chain polyunsaturated ω3 fatty acids (VLCPFA), docosahexaenoic acid (DHA, 22:6) and eicosapentaenoic acid (EPA, 20:5). DHA had the highest amount in both cultures (Figure 4), so we used it as a standard to estimate relative abundance of other fatty acids (Table 2). Trace amount of tetracosahexaenoic acid, a C24 VLCPFA, was found. Three saturated fatty acids, palmitic acid (16:0), stearic acid (18:0), and behenic acid (20:0), were also identified. The abundance of saturated fatty acids was relatively low in the phosphate-depleted cultures. The analysis also revealed the presence of C16 unsaturated fatty acids: 7-hexadeceonoid acid (16:1), 9-hexadecenoid acid (16:1), 7,10-hexadecadienoic acid (16:2), 7, 10, 13-hecadecatrienoic acid (16:3), and 4, 7, 10, 13-hexadecatetraenoic acid (16:4) in both of the cultures. In the phosphate-depleted cultures, the relative abundance of C16 and C18 fatty acids was lower than the fatty acids in the phosphate-replete cultures (Figure 4 and Table 2), suggesting that phosphate availability in the culture medium affected the lipid composition in *Eutreptiella* sp.

**Figure 4.**
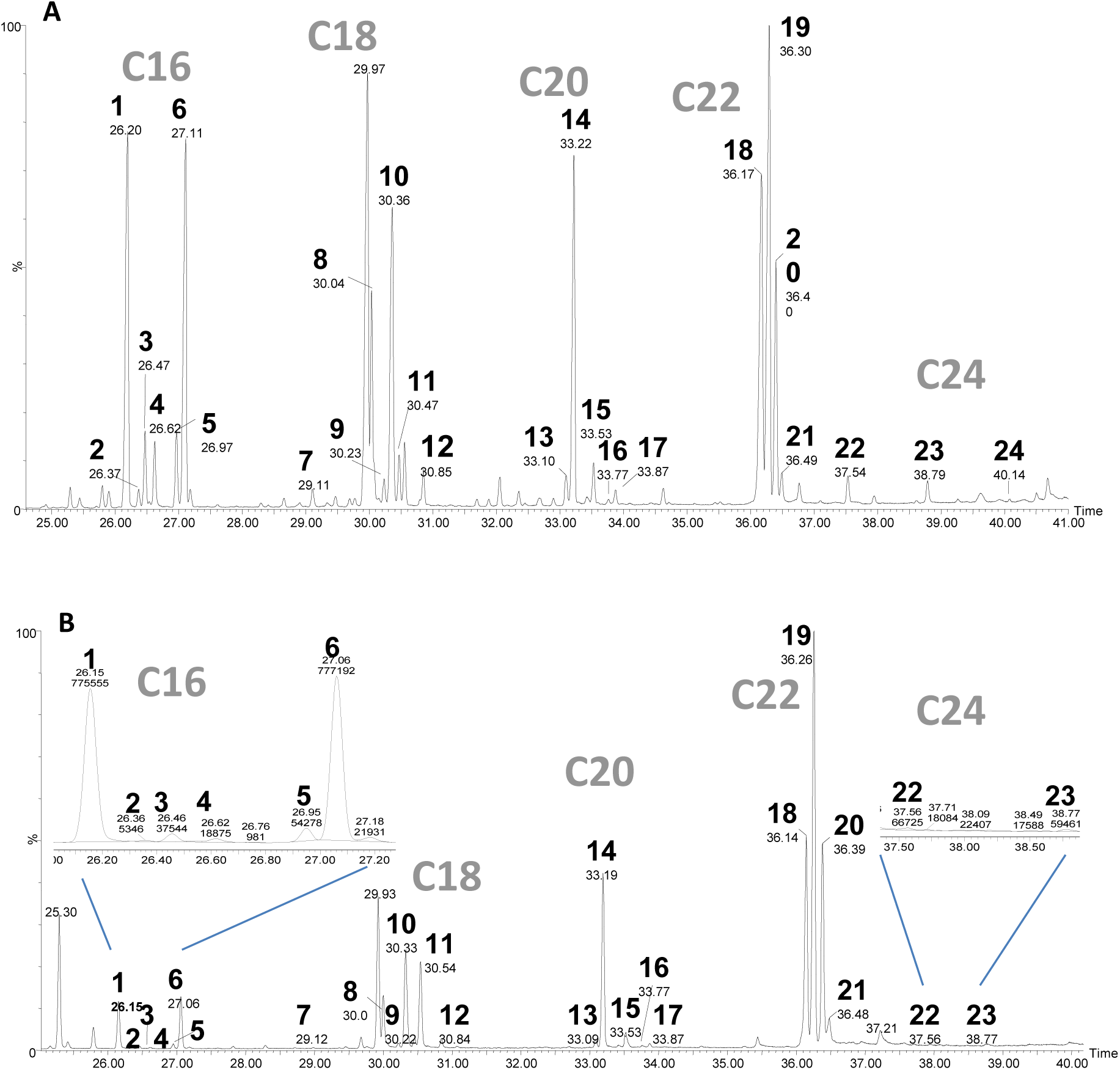
Chromatograms of total fatty acid esters from *Eutreptiella* sp. Note that DHA (peak 19) was the most abundant fatty acid. Corresponding fatty acids (labeled by length of carbon, e.g. C16) were listed in Table 3. (A) Chromatogram of fatty acid esters from the phosphate–replete cultures. (B) Chromatogram of fatty acid esters from the phosphate-depleted cultures. Insets are closer-ups of some of the small peaks of esters. The first number (bold-typed) by the peak is peak ID, the second number retention time, and the third number peak area (only trace elements are shown). Note that retention times between the two samples are slightly different.

Based on the transcriptome and GC/MS/MS results, *Eutreptiella* sp. is predicted to be capable of producing various intermediates via ω3 and ω6 pathways, as well as producing VLCPFA EPA and DHA by using the Δ8 alternative pathway, as reported in *Euglena* (Meyer et al. 2003; Qi et al. 2004; Wallis and Browse 1999). Figure 5 shows the potential fatty acid elongation pathways for *Eutreptiella* sp. as constructed from the transcriptome results. The first desaturation for unsaturated fatty acids synthesis is catalyzed by Δ9 desaturase to introduce a double bond into stearic acid resulting in 18:1 –ACP. Then a series of elongation and desaturation takes place to produce the 24 types of unsaturated fatty acid mentioned before (Figure 5). The elongation procedure is terminated when the acyl group is removed by acyl-ACP thioesterase (TE) or oleoyl-ACP hydrolase (OAH), or when the fatty acids are transferred to the biosynthesis of triacylglycerol.

**Figure 5.**
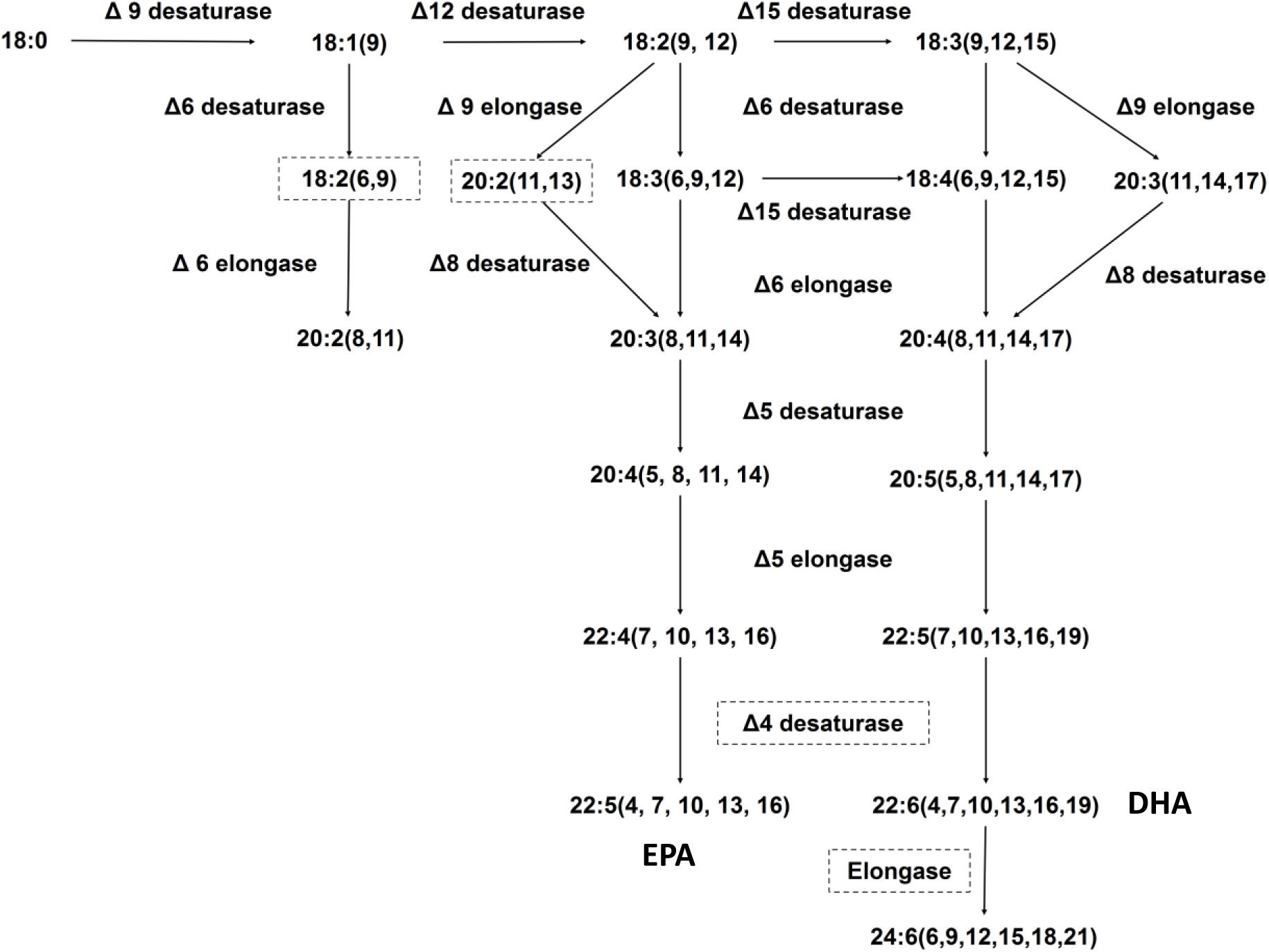
Putative pathways of fatty acids elongation and desaturation inferred from *Eutreptiella* sp. transcriptome annotation and GC/MS/MS analyses. Boxes with dashed lines indicate the enzymes or substrates that were not found in our study.

### Triacylglycerol (TAG) biosynthesis

Microalgal TAG biosynthesis is proposed to take place via transfer of fatty acids from CoA to glycerol-3-phosphate using the direct glycerol pathway (Ratledge 1988). However, the existing knowledge on the pathways and enzymes involved in TAG biosynthesis in most of the microalgae (e.g. euglenophytes) are limited. In this study, four transcripts coding for the enzymes involved in TAG biosynthesis were found in our *Eutreptiella* sp. transcriptomic dataset. They were glycerol 3-phosphate dehydrogenase (GPD), glycerol 3-phsphate acyltransferase (GPAT), and acylglycerophosphate acyltransferase (AGPAT) (Table 1). For TAG biosynthesis, the precursor glycerol 3-phosphate is produced during glycolysis from dihydroxyacetone phosphate by the action of glycerol 3-phosphate dehydrogenase.

Based on the data, a potential pathway for *Eutreptiella* sp. TAG biosynthesis is proposed (Figure 3). The first step is the acylation of the two hydroxyl groups of glycerol3-phosphate by two fatty acyl-CoA to yield phosphatidic acid, which is hydrolyzed by phosphatidate phosphatase to form 1,2 diacylglycerol. Diacylglycerols are then converted to triacylglycerols by transferring a third fatty acyl-CoA. Phosphatidic acid phosphohydrolase (PAP) and diacylglycerol acyltransferase (DGAT) were missing in our cDNA libraries.

### Gene expression levels as measured using reverse transcription quantitative PCR

We further confirmed the differential expression patterns of several genes using reverse transcription quantitative PCR (RT-qPCR). Nine genes involved in lipid synthesis were chosen for the analysis, including type I fatty acid synthase (FASI), 3 different types of β –ketoacyl-ACP reductase (KAR1, KAR2, KAR3, 12-33% identity at amino acid level), acetyl-CoA carboxylase (ACCase), acyl carrier protein (ACP), thioesterase (TE), Δ8 desaturase, and an unclassified fatty acid desaturase (FAD). Seven of the nine genes, KAR2, KAR3, FAD, ACCase, ACP, Δ8, and TE had higher expression levels in the high-lipid-light cultures (Figure 6 and Figure 7). The expression of FASI showed an opposite trend between high-lipid and low-lipid cultures. In high-lipid cultures, FASI was expressed at a higher level in the light period, whereas in the low-lipid cultures, FASI was expressed at a lower level in the light period (Figure 6). As for genes involved in triacylglycerol synthesis, glycerol 3-phosphate dehydrogenase (GPD) and 1-acyl-sn-glycero3-phosphate o-acyltransferase (AGPAT) exhibited slightly higher expression (transcript abundance) in high-lipid-light cultures (Figure 7).

**Figure 6.**
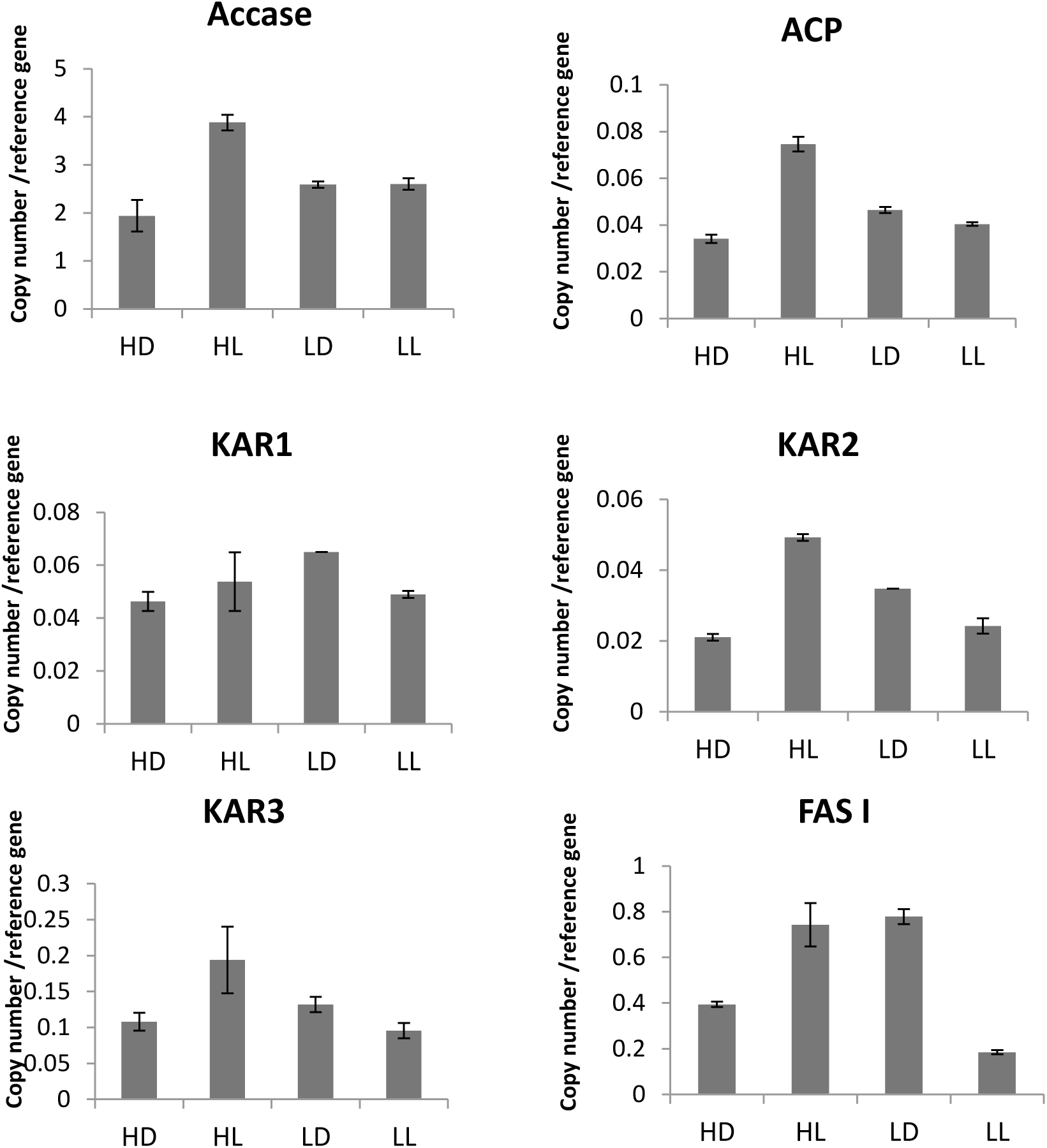
Expression levels of genes involved in fatty acids biosynthesis in *Eutreptiella* sp. HD: high-lipid-dark cultures. HL: high-lipid-light cultures. LL: low-lipid-light cultures. LD: low-lipid-dark cultures.

**Figure 7.**
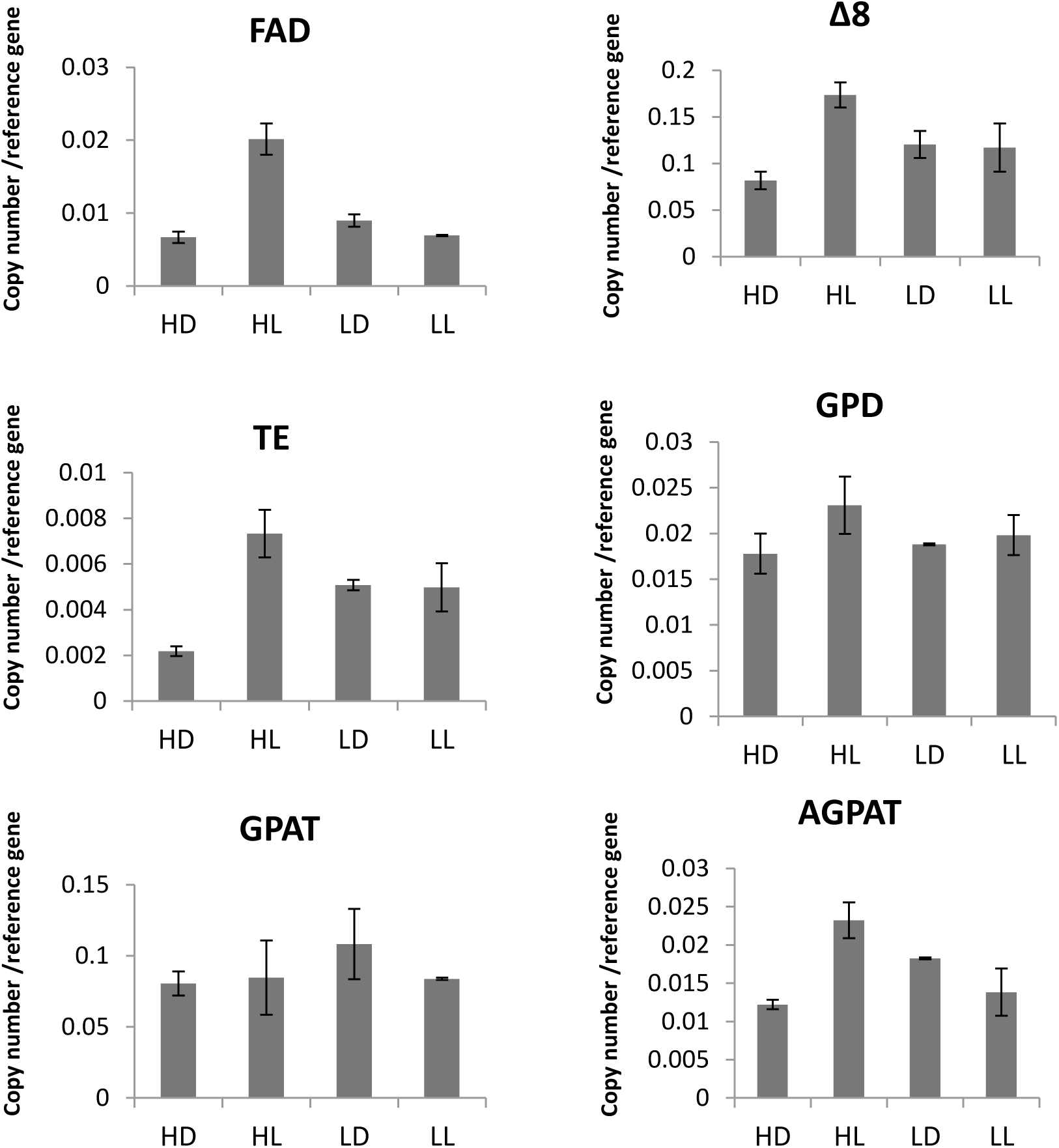
Expression levels of genes involved in fatty acids biosynthesis (FAD, Δ 8, and TE) and triacylglycerol biosynthesis (GPD, GPAT, and AGPT). HD: high-lipid-dark cultures. HL: high-lipid-light cultures. LL: low-lipid-light cultures. LD: low-lipid-dark cultures.

We also examined the expression of genes potentially involved in carbon fixation in *Eutreptiella* sp.: phosphoenolpyruvate carboxylase (PEPCase), 2 variants of phosphoenolpyruvate carboxykinase (PEPCK1 and PEPCK2, 54% identical at amino acid level), pyruvate-phosphate dikinase (PPDK), and phosphoribulokinase (PRK). Three out of the five genes (i.e. PPDK, PRK, and PEPCK1) showed positive correlation of transcript abundance with lipid production. In particular, PEPCK1 and PRK expression levels in the high-lipid-light cultures were over 6- and 4-fold higher than in the low-lipid-light cultures, respectively (Figure 8).

**Figure 8.**
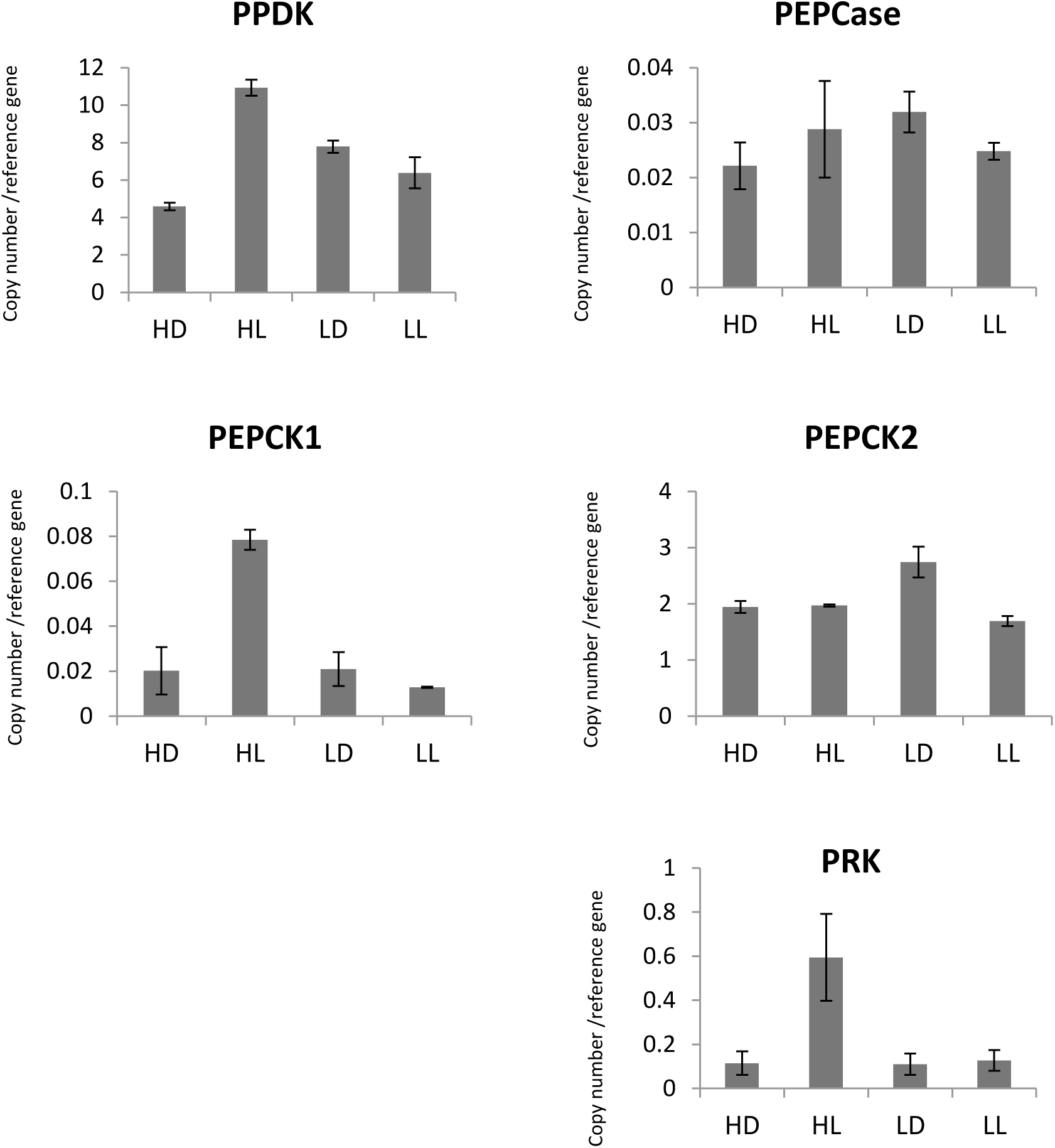
Expression levels of genes involved in carbon fixation. HD: high-lipid-dark cultures. HL: high-lipid-light cultures. LL: low-lipid-light cultures. LD: low-lipid-dark cultures.

## Discussion

### Unique lipid and fatty acid production patterns

While the effects of depleting nitrate on lipid production has been studied in many algae (Griffiths and Harrison 2009, Hu et al. 2008), effects of phosphate are less documented. Our preliminary experiments showed that phosphate depletion more strongly promotes lipid content in *Eutreptiella* sp. cells than nitrate depletion (data not shown). The study reported here also showed that phosphate stress markedly increased *Eutreptiella* lipid content. Among the cultures grown in parallel, lipid content was 2 fold as much in the phosphate-depleted cultures as that in the phosphate-replete, f/2 medium-grown, cultures (Figure 1). The type of main energy reserve (carbohydrates or lipids) in algae depends on which carbon fixation product is utilized in the Calvin-Benson cycle. Among those products, hexose phosphates are used in the synthesis of carbohydrates, while 3-phophoglycerate is used for lipid synthesis (Raven 1974). In addition, the availability of inorganic nitrogen influences carbon partition (Huppe and Turpin 1994, Raven 1974). The presence of inorganic nitrogen source favors 3-phosphoglycerate and increases the flux of photosynthetically fixed carbon into citrate cycle (Raven 1974), which is one of the sources for acetyl CoA (Nelson and Cox 2008).

Interestingly, *Eutreptiella* sp. has abundant polyunsaturated fatty acids. DHA was the most abundant fatty acid from both the phosphate-depleted and phosphate-replete cultures. Polyunsaturated fatty acids are valuable for cold adaptation (Wallis et al. 2002 and references therein). The relatively high abundance of unsaturated fatty acids in *Eutreptiella* sp. may be due to the adaptation to their native environment in Long Island Sound, where the temperature reaches down below in the winter although there has been a warming trend (Snyder et al. 2019). All the fatty acids that we identified in this study have also been reported in *Euglena gracilis* (Hulanicka et al. 1964, Korn 1964, Meyer et al. 2003), except C24:1 and C24:6 fatty acids. Different from *E. gracilis, Eutreptiella* sp. has a very high proportion of very long chain unsaturated fatty acids (C22 and longer, i.e. DHA and EPA), rendering this species to be of high nutritional value for zooplankton and other heterotrophic organisms in the marine ecosystem. Moreover, the proportions of saturated fatty acids (i.e. C16:0, C18:0, and C20:0) and unsaturated C16 and C18 fatty acids decreased when phosphate was depleted (Figure 4 and Table 2), suggesting that it is possible to enhance the production of desired components of fatty acids from *Eutreptiella* sp. by manipulating nutrient composition in the culture medium. Although it is not clear why the abundance of C16:0 and C16:4 fatty acids were much lower in the phosphate-depleted cultures, phosphate stress might alter the composition of fatty acids in the algal cells, as plants and algae can respond to stress by adjusting membrane fluidity and by releasing unsaturated fatty acids from the membrane (Maréchal et al. 1997, Upchurch 2008).

### Expressed genes linked to lipid synthesis in *Eutreptiella* sp

Fatty acids are synthesized via two enzyme systems, ACCase and FAS (Nelson and Cox 2008). ACCase plays a critical role in regulating fatty acid synthesis and is essential for lipid metabolism as it catalyzes the first step of fatty acid synthesis and is found in all kingdoms of life except archaea (Cronan Jr and Waldrop 2002, Harwood 1988). It comprises three functional constituents: biotin carrier protein, biotin carboxylase, and transcarboxylase, and differentiates into two distinct types: one consisting of three separate subunits (prokaryote-type), and the other being a single multi-domain multifunctional polypeptide (eukaryote-type) (Sasaki et al. 1995). The eukaryote-type occurs in mammals, fungi or yeast, while both the prokaryote- and the eukaryote-type can be found in plants (Huang et al. 2002, Sasaki et al. 1995). The ACCase sequences found in our dataset shares ∼72% amino acid identity to an eukaryotic type in the diatom *Thalassiosira pseudonana*, suggesting that *Eutreptiella* sp. ACCase is also an eukaryotic type. In the qPCR analysis, ACCase showed highest expression level in the high-lipid-light cultures, indicating that this gene is related to photosynthetic lipid production in *Eutreptiella* sp.

Our results suggest that *Eutreptiella* sp. may have both type I fatty acid synthase (FASI) and type II fatty acid synthase (FASII), which are the two major types of fatty acid synthases that naturally occur in different groups of organisms (Nelson and Cox 2008). FASI, found in vertebrate and fungi, is a single multifunctional polypeptide with 7 active sites, whereas FASII, found in plants and bacteria, compose discrete and freely diffusible enzymes. Interestingly, *Euglena gracilis*, which is another euglenoid species, has different types of fatty acid synthesis pathways and enzymes to synthesize lipids under different environmental conditions (Delo et al. 1971, Goldberg and Bloch 1972, Inui et al. 1984). ACP-dependent FASI is located in the cytosol and mainly produces C16 fatty acids (Goldberg and Bloch 1972). When grown in the light, *E. gracilis* utilizes ACP-dependent FAS II in plastids to produce fatty acids (Ernst-Fonberg and Bloch 1971). Inui et al. (1984) also reported a malonyl-CoA independent fatty acid synthesis system to form wax ester from paramylon as energy storages in *E. gracilis* under anaerobic conditions. We found genes coding for FASI and for β –ketoacyl-ACP reductase and Acyl-ACP thioesterase of FASII in *Eutreptiella* sp. Our failure to retrieve the rest of the FASII gene complex from our cDNA libraries could be due to their being plastid-coded genes, which would not contain Eut-SL that we used to construct the cDNA libraries, or just simply because they were not expressed. Whether *Eutreptiella* sp. contains the other type of ACP-dependent FAS similar to that in *E. gracilis* remains to be further investigated in the future.

Based on the fatty acid composition and possession of genes coding for enzymes involved in fatty acid elongation and desaturation in *Eutreptiella* sp., it is evident that the ω3, ω6 and alternative Δ8 pathways occur in the euglenoid algae. Different from most of plants and most algae, we found that *Eutreptiella* is similar to *Euglena* in using Δ8 desaturase to introduce the third double bond on 20:2 (11, 14) to form 20:3 (8, 11, 14) (Meyer et al. 2003, Qi et al. 2004). The 20:3 fatty acid is then converted to 22:5 EPA via stepwise Δ5 desaturase, Δ5 elongase and Δ4 desaturase (Figure 5). In order to produce DHA, *Eutreptiella* may also utilize another Δ8 pathway to introduce the 4^th^ double bound on 20:3 (11,14,17), resulting in 20:4 (8,11,14,17) for later stepwise process of elongation and desaturation. Although Δ4 desaturase was not identified in our dataset, it is very likely that *Eutreptiella* sp. has this gene since it was identified and involved in DHA synthesis in *E. gracilis* (Meyer et al. 2003). In our study, both EPA would need Δ4 desaturase to form their last double bond. We also detected trace amount of 15-tetracosenoic acid (24:1) and 6,9,12,15,18,21-tetracosahexaenoic acid (24:6). The 24:6 fatty acid can be found in some algae (Mansour et al. 2005), but the synthesis pathway is not well studied. We proposed that the 24:6 fatty acid could potentially be synthesized by elongation from DHA or elongation and desaturation from other C22 fatty acids.

The 16:4 (4,7,10,13) fatty acid was also abundant in phosphate-replete cultures of *Eutreptiella* sp. Highly unsaturated stearidonic acid (18:4) and hexadecatetraenoic acid (16:4) are produced in some macroalgae (Dembitsky et al. 1991, Ishihara et al. 2000) and microalgae (Wallis and Browse 1999). In diatoms, the phytoplankton accounting for about 40% of algal primary production, the most common fatty acids range from C14:0 to C22:6, with the most common number of double bonds two to three and rarely more than 6 (Yi et al. 2017). Unsaturated C16 fatty acids have been identified in *E. gracilis*, the close relative of *Eutreptiella* sp. (Constantopoulos and Bloch 1967). Increasing light intensity can enhance the production of 16:4 acid and 18:3 α-linolenic acid in *E. gracilis* (Constantopoulos and Bloch 1967). Korn (1964) suggested a possible C16 pathway for *E. gracilis*: 16:1 (7) → 16:2 (7, 10) → 16:3 (7, 10, 13) →16:4 (4, 7, 10, 13). Although we identified 16:4 (4, 7, 10, 13) in *Eutreptiella* sp., the genes responsible for the synthesis of this abundant fatty acid remain uncertain.

### Regulation of genes associated with lipid production

Our RT-qPCR revealed that many of the genes examined had different expression levels between the dark and light periods. Genes involved in high lipid biosynthesis showed higher expression levels under light condition, suggesting that these genes promoted lipid production in *Eutreptiella* sp. during photosynthesis. Those genes were ACCase, ACP, KAR2, KAR3, FAD, Δ8 desaturase, TE, AGPAT, and GPD. The results of RT-qPCR verified our physiological observation from a 24h diel experiment, showing that lipids in *Eutreptiella* sp. cells start to accumulate during the light period after cells have actively divided (Kuo and Lin, 2013). Other evidence showed that the activity of ACCase is correlated to that of the Calvin-Benson Cycle enzymes, with the greatest activities under light (Sasaki et al. 1997) and negligible in the dark (Bao et al. 2000).

Three genes encoding carbon fixation enzymes showed higher expression levels under the high-lipid-light condition, indicating that those enzymes are related to the biosynthesis of precursors for lipid production. Phosphoribulokinase (PRK) and one copy of phosphoenolpyruvate carboxykinase (i.e. PEPCK1) exhibited the highest level of relative gene expression among these genes. PEPCK activity is linked to CO_2_ fixation in algae (Reiskind and Bowes 1991). It converts oxaloacetate into CO_2_ and phosphoenolpyruvate, bridging C4 to C3 pathway in providing CO_2_ for photosynthesis. It is noteworthy that C4 enzymes have been identified in this species (Kuo et al. 2013). The phosphoenolpyruvate can later be converted into pyruvate, which is an important precursor for Acetyl-CoA. PRK catalyzes the reaction of ATP and ribulose 5-phosphate to produce ADP and ribulose 1,5-bisphosphate, the substrate needed to capture CO_2_ in photosynthesis. The concomitant up-regulation of the fatty acid synthesis and photosynthesis genes suggests the linkage of the two carbon metabolic processes.

## Supporting information

Table S1. Lipid biosynthesis genes

## Acknowledgements

We thank the Center of Environmental Science and Engineering at UConn for providing the facility of GC/MS/MS, and Yunyun Zhuang for helping with 454 sequencing. The research was supported by the Predoctoral Awards of the Department of Marine Sciences at UConn (to RCK), Multidisciplinary Environmental Research Award from the Center of Environmental Science and Engineering at UConn (to RCK), and National Science Foundation “Assembling the Tree of Life (AToL)” grant EF-0629624 (to SL and collaborators). All the authors declare no conflict of interest existing in this work.

## Notes

### Competing Interest Statement

The authors have declared no competing interest.

### Summary of Updates

Correct English grammatical errors and rephrased a few sentences.

